# *Xist* expression in male *Peromyscus leucopus* is associated with restricted chromatin repression and incomplete X-to-autosome dosage compensation

**DOI:** 10.64898/2026.06.11.731719

**Authors:** Maria De Lourdes Andrade Ludena, Jonathan V. Duong, Aqsa Motiwala, Christopher L Hartl, Anthony D. Long, Alan G. Barbour, Sha Sun

**Affiliations:** Department of Developmental and Cell Biology, Charlie Dunlop School of Biological Sciences, University of California, Irvine, Irvine, CA, USA; Department of Microbiology and Molecular Genetics, School of Medicine, University of California, Irvine, Irvine, CA, USA; Epigenome Technologies, San Diego, CA, USA; Department of Systems Biology, Charlie Dunlop School of Biological Sciences, University of California, Irvine, Irvine, CA, USA; Department of Medicine, School of Medicine, University of California, Irvine, Irvine, CA, USA

**Keywords:** X-chromosome inactivation, Xist, dosage compensation, sex chromosome, long noncoding RNA

## Abstract

X-chromosome inactivation (XCI) equalizes X-linked gene dosage between XX females and XY males in eutherian mammals and is canonically initiated by the long noncoding RNA *Xist*, which coats one X chromosome and recruits Polycomb-mediated chromatin silencing. *Xist* expression has therefore been considered female-specific and central to balancing gene dosage between two X chromosomes in females and a single X chromosome in males. However, emerging evidence suggests that *Xist* can also be expressed in male cells in both healthy and pathogenic contexts, raising fundamental questions about the scope and constraints of *Xist* function. Here, we identify robust *Xist* expression in somatic cells of male white-footed deermice (*Peromyscus leucopus*), a placental mammal with a conventional XY karyotype. Unlike the chromosome-wide *Xist* RNA coating observed in females, male *Xist* RNA localizes as discrete nuclear puncta and is detected across multiple tissues by RNA-seq, RT-qPCR, and RNA fluorescence *in situ* hybridization. Integrative transcriptomic and chromatin profiling using CUT&Tag and single-nucleus Paired-Tag reveals that male *Xist* expression is not merely incidental but has measurable regulatory consequences. In females, *Xist* expression is associated with chromosome-wide enrichment of the repressive histone mark H3K27me3 and transcriptional silencing of X-linked genes. In males, although *Xist* transcription is also associated with H3K27me3, this interaction does not induce global X-chromosome inactivation. Instead, *Xist* selectively represses a subset of dosage-sensitive neural and mitochondrial X-linked genes, while the majority of X-linked genes remain largely active. Notably, with persistent *Xist* activity, neither males nor females achieve complete X-to-autosome dosage compensation in *P. leucopus*. These findings establish *P. leucopus* as a mammalian system in which male *Xist* operates outside the canonical female-specific XCI pathway, revealing a gene-selective model of X-chromosome regulation. Our results challenge the universality of prevailing XCI models and suggest that partial, context-dependent *Xist* activity represents a viable and evolutionarily stable dosage compensation strategy in mammals.

**Author Summary:** In mammals, females and males differ in their sex chromosome: females have two X chromosomes, while males have one. This difference creates a potential imbalance in gene dosage, which can disrupt development if not properly controlled. Typically, females compensate by turning off one chromosome through a process called X-chromosome inactivation (XCI), which is triggered by the long noncoding RNA *Xist*. In parallel, a process known as X-to-autosome dosage compensation balances X-linked gene expression relative to the rest of the genome. Because XCI occurs in females, *Xist* has long been considered to function only in female cells and be inactive in males. Here, we show that *Xist* is persistently expressed in male white-footed deermice (*Peromyscus leucopus*), a cricetid rodent with a conventional XY sex chromosome system. Unlike the large *Xist* RNA “cloud” that covers the inactive X chromosome in females, *Xist* RNA in males appears as small, discrete spots in the nucleus. By combining gene expression and chromatin profiling, we find that male *Xist* associates with the repressive histone mark H3K27me3 but does not switch off the entire X chromosome. Instead, *Xist* in males selectively reduces the activity of a limited set of X-linked genes, while most X-linked genes remain active. Surprisingly, we also find that neither male nor female *P. leucopus* achieves complete balance between X-linked and autosomal gene expression, despite the long-held view that such balance is essential for mammalian viability. By uncovering functional *Xist* activity in males of a non-model species and demonstrating incomplete dosage compensation in both sexes, this work challenges the traditional view of XCI as a strictly female-specific strategy and reveals an unexpected flexibility in how mammals regulate the X chromosome.

## Introduction

The XX female and XY male sex chromosome system in eutherian mammals evolved approximately 170 million years ago from a pair of autosomes (1,2). Acquisition of a male sex-determining gene led to suppression of recombination between the proto-X and proto-Y chromosomes; and as the Y chromosome subsequently degenerated, males were left with a single copy of hundreds of genes that remain duplicated in females. Placental mammals resolved this imbalance through X-chromosome inactivation (XCI), which transcriptionally silences one X chromosome in each female somatic cell (3–5). XCI is mediated by *X-inactive specific transcript* (*Xist*), a long noncoding RNA (lncRNA) expressed exclusively from the inactivate X chromosome (6,7). In mice and humans, *Xist* spreads *in cis* across the X chromosome and recruits repressive chromatin machinery to establish gene silencing (8,9). Proper XCI and dosage compensation are essential for early development: complete failure results in embryonic lethality, whereas incomplete XCI leads to developmental defects associated with dysregulation of X-linked genes (4,10,11).

Studies in mouse and human models have shown that the core mechanisms of XCI, initiated by the *Xist* lncRNA, are largely conserved. The *Xist* transcript contains multiple conserved domains, including tandem repeats, Repeat A, B, C, D, E, F, which recruit transcriptionally repressors required for its function. Among these, Repeat A is the most conserved and essential repeat, mediating recruitment of Polycomb repressive complex 2 (PRC2) to the inactive X chromosome, leading to the deposition of the repressive histone mark H3K27me3 (histone 3 lysine 27 trimethylation), a hallmark of X chromosome silencing (12–14). Disruption of this pathway is typically incompatible with development, and ectopic *Xist* expression in male cells, which carry a single X chromosome, has been considered lethal due to silencing of essential X-linked genes (8,14). Accordingly, *Xist* has been classically viewed as a female-specific RNA whose function is intrinsically linked to chromosome-wide dosage compensation. However, emerging evidence challenges this paradigm. *XIST* expression has been detected in male testicular germ cell tumors, and pan-cancer analyses reveal somatic activation of *XIST* across diverse male malignancies, often accompanied by partial X-chromosome silencing, altered chromatin accessibility, and changes in DNA methylation (15). Furthermore, experimental induction of *Xist* in male mice triggers immune dysregulation and lupus-like phenotypes, while male patients with systemic lupus erythematosus exhibit elevated *XIST* expression and partial repression of X-linked gene repression (16,17). Together, these findings demonstrate that male cells can engage components of the XCI machinery, but whether this reflects pathological deregulation or a broader, context-dependent capacity for *Xist* function remains unresolved.

The current model of *Xist* regulation is based largely on studies of model organisms, particularly the house mouse *Mus musculus*. However, species-specific variations in XCI regulation and noncanonical *Xist* functions have been increasingly recognized (18,19). Despite *Xist*-mediated silencing along the inactive X chromosome, a subset of X-linked genes escape inactivation and remains expressed from both the active and inactive X chromosomes (20). This escape from XCI occurs in approximately 3–7% of X-linked genes in mice and 12–20% in humans (21–23). These observations highlight variability in *Xist* function and its silencing efficiency, underscoring the need for broader comparative studies to better understand the role *Xist* and the evolution of dosage compensation mechanisms in mammals.

Advances in genome sequencing of the white-footed deermouse (*Peromyscus leucopus*) provide new opportunities to investigate chromosomal regulation and XCI in a non-model organism (24). Unlike commonly studied murine models such as *Mus musculus* (house mouse) and *Rattus norvegicus* (brown rat), *Peromyscus* species belong to the Family Cricetidae and are evolutionarily closer to hamsters and voles than to the Family Murinae including rats and house mice (Fig. 1A). Some cricetids exhibit diverse and unconventional patterns of sex chromosome composition and X chromosome regulation. For example, males of the creeping vole (*Microtus oregoni*) carry two X chromosomes and show male-specific *Xist* expression (25,26). In the mole vole genus (*Ellobius*), the Y chromosome is absent, and both sexes exhibit XO or XX karyotypes, raising fundamental questions about how dosage compensation is adapted (27). Wood lemmings (*Myopus schisticolor*) further exemplify this diversity, displaying sex-specific and mosaic XCI patterns associated with a variant X* chromosome – a modified X chromosome that can feminize XY individuals and alter XCI dynamics (28).

**Figure 1.**
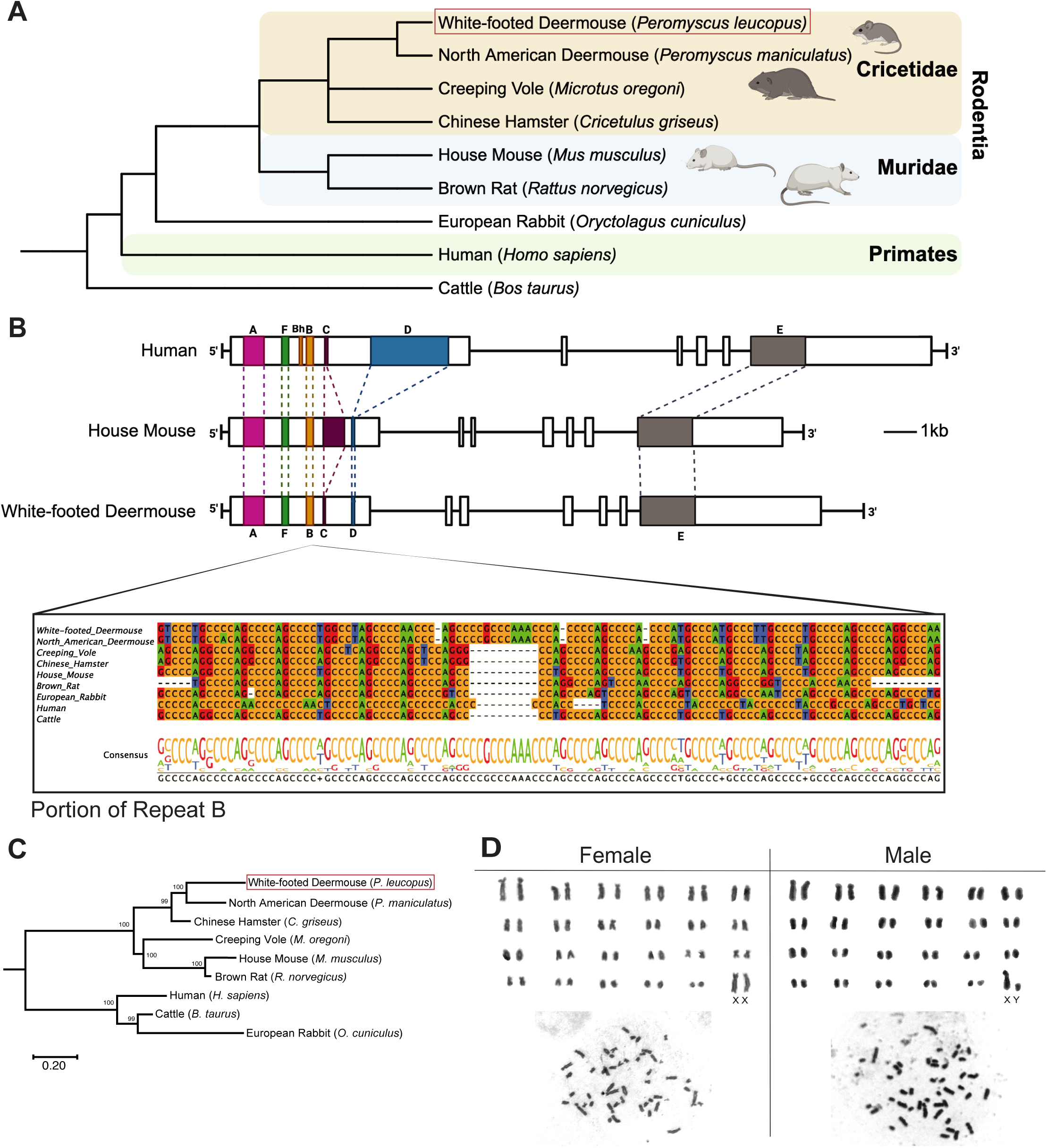
Phylogenetic analysis and sequence comparison of *Xist* in eutherian species. (A) Whole-genome cladogram of nine representative eutherian mammals illustrating the divergence between Muridae (blue) and Cricetidae (tan) within Rodentia. *Peromyscus leucopus* and *P. maniculatus* cluster within Cricetidae, whereas *Mus musculus* and *Rattus norvegicus* represent Muridae. *Oryctolagus cuniculus*, *Homo sapiens*, and *Bos taurus* serve as outgroups. (B) Gene structure of *Xist* in human, mouse, and *P. leucopus*, with conserved repeat domains (Repeat A–F) aligned at the exon level. A multiple sequence alignment of a representative region from Repeat B illustrates conservation of the Polycomb-interacting domain. (C) Maximum-likelihood phylogeny based on *Xist* nucleotide sequences recapitulates known evolutionary relationships and suggests a close relationship of *Microtus oregoni* to murid species. Node labels indicate percent bootstrap support. (D) Karyotype analysis of *P. leucopus* dermal fibroblasts confirms a diploid chromosome count (2n = 48) in both sexes. Shown are validated karyotypes from a female (XX) and a male (XY) individual.

To investigate how *Xist* function may diverge in different evolutionary contexts, we examine its activity in *Peromyscus leucopus*, a non-murine rodent that provides a phylogenetically distinct contrast to canonical model systems. Using an integrative approach combining comparative sequence analysis, single-cell imaging, transcriptomics, and chromatin profiling, we define the regulatory consequences of *Xist* expression in *P. leucopus* and test whether *Xist* function is intrinsically coupled to chromosome-wide XCI and dosage compensation. We find that *Xist* is expressed in both sexes and recruits Polycomb-associated repressive chromatin in a context-dependent manner. However, rather than inducing chromosome-wide silencing, *Xist* selectively represses a subset of X-linked genes, while most remain transcriptionally active in *P. leucopus* males. Consistent with this, *P. leucopus* fails to achieve X-to-autosome dosage compensation, the balancing of X-linked gene expression relative to autosomes.

## Results

### *Xist* Sequence Domains are Identified in the *Peromyscus leucopus* Genome

To investigate how the long noncoding RNA *Xist* has evolved across placental mammals, we compared its sequence and structure features in nine eutherian species, including rodents and humans (Fig. 1). Although *Xist* plays a conserved role in X-chromosome silencing, its nucleotide sequence as a noncoding RNA shows substantial divergence among species. For example, *Xist* (house mouse) and *XIST* (human) share an overall sequence identity of 49%, which is markedly lower than the ∼85% identity typically observed for protein-coding genes between the mouse and human genomes, which diverged approximately 90 million years ago (29,30).

We next analyzed the *Xist* locus in the whole-genome assembly of white-footed deermouse (*Peromyscus leucopus*), which diverged from the house mouse (*Mus musculus*) 25-40 million years (24). Despite overall sequence divergence, we focused on whether key functional elements, repetitive domains known as Repeats A–F, are conserved (12,31,32). Comparative analysis of syntenic X-chromosome regions of human, house mouse, and deermouse revealed that the overall *Xist*/*XIST* architectural organization is conserved (Fig. 1B). Notably, while *P. leucopus Xist* contains a minimal version of the C and D repeats, it retains a C-rich B repeat, which is critical for the Polycomb Interaction Domain (PID). This domain, comprising the entire B repeat and part of the C repeat, is required for recruitment of Polycomb complexes that deposit repressive histone marks such as H3K27me3 (32). The identification of repetitive domains suggests that *Peromyscus Xist* preserves key functional motifs necessary for the X-chromosome silencing capacity, supporting functional conservation despite sequence divergence typical of lncRNAs.

Phylogenetic analysis of full-length *Xist* sequences across nine eutherian species recapitulated expected rodent taxonomy, with murids (*M. musculus*, *R. norvegicus*) separating from cricetids including *Peromyscus, O. torridus, and,C. griseus* (Fig. 1C; Supplementary Fig. 1). Interestingly, *M. oregoni*, a cricetid species with XX males, clusters more closely with murids than with other cricetids. This unexpected placement, deviating from the whole-genome phylogeny, is likely driven by conservation of the C repeat in *M. oregoni,* which resembles the murid C repeat in size and structure.

To confirm that subsequent functional analyses were conducted in a conventional XX/XY sex chromosome context, we verified the karyotype of *P. leucopus*. Cytogenetic analysis of primary dermal fibroblasts confirmed the expected diploid complement (2n = 48) (Fig. 1D), with females carrying two X chromosomes (XX) and males carrying one X and one Y chromosome (XY), consistent with prior reports (33–36).

### *Xist* Expression is Detected in *P. leucopus* Dermal Fibroblasts

To investigate whether *Xist* is transcriptionally active and functionally engaged in *P. leucopus*, we examined its gene expression and subnuclear localization in dermal fibroblasts from both *P. leucopus* and *M. musculus*. These primary cell cultures comprise heterogeneous dermal fibroblast cells, including at least two transcriptionally distinct subtypes (37), allowing assessment of *Xist* activity across cellular states. Using species-specific sequence primers targeting *Xist* exon 1, RT-PCR (RNA reverse transcription followed by PCR) detected robust *Xist* expression in female dermal fibroblasts from both species, consistent with canonical *Xist* activity and XCI. As expected, male *M. musculus* fibroblasts exhibited minimal *Xist* expression. In contrast, male *P. leucopus* dermal fibroblasts consistently produced a distinct *Xist* amplicon (Fig. 2A), indicating that *Xist* transcription is not restricted to females in this species.

**Figure 2.**
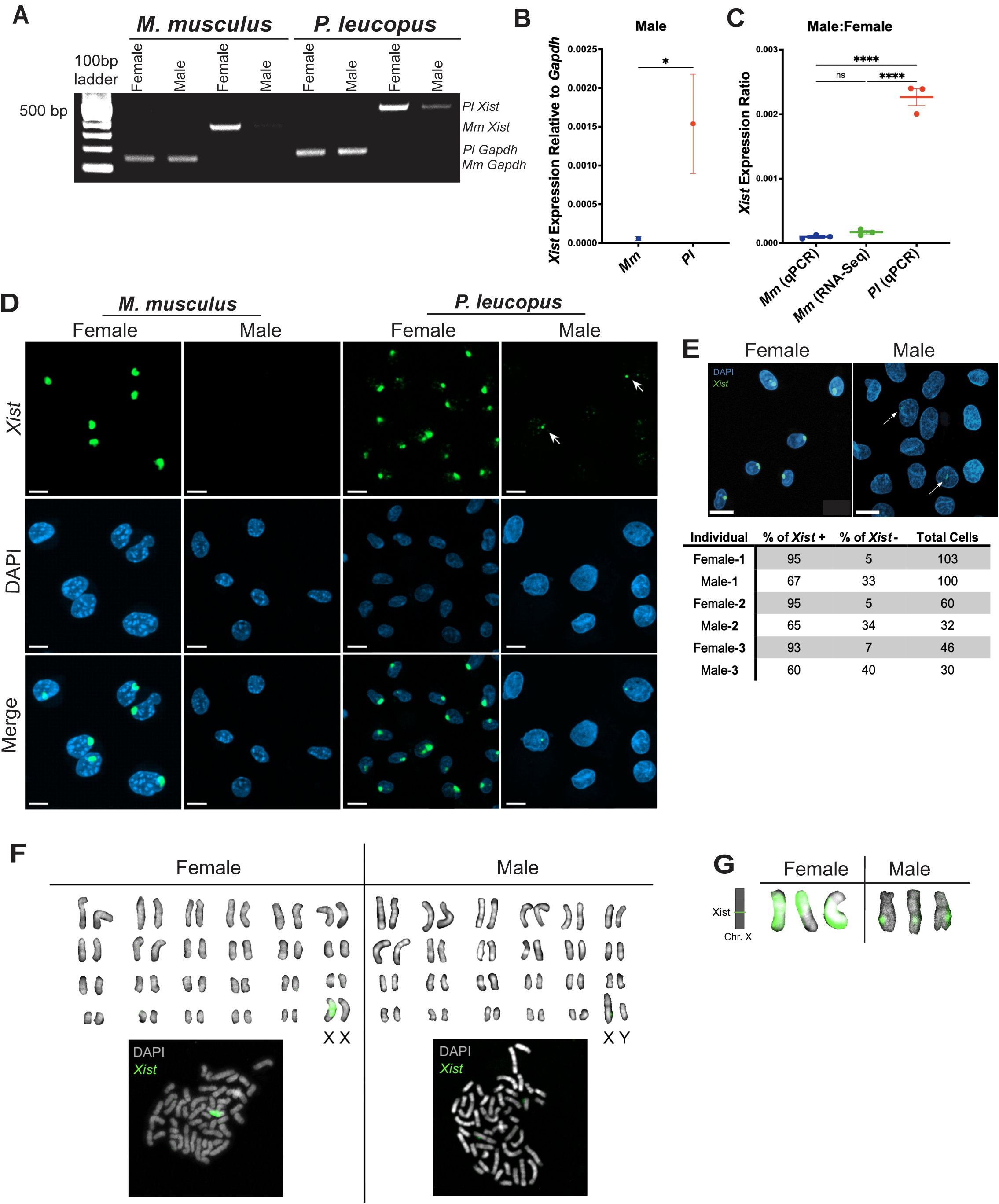
*Xist* expression in *M. musculus* and *P. leucopus* primary dermal fibroblasts. (A) RT-PCR demonstrates robust *Xist* expression in female fibroblasts of both species and detectable *Xist* transcript in male *P. leucopus* but not *M. musculus*. *Gapdh* serves as a control for cDNA quality. (B) RT-qPCR quantification shows significantly higher relative *Xist* expression in male *P. leucopus* fibroblasts compared with *M. musculus* (n = 3 biological replicates per sex; mean ± SEM; *p* < 0.05, t-test). (C) Male-to-female *Xist* expression ratios in *M. musculus* fibroblasts (qPCR, this study), multiple *M. musculus* tissues (bulk RNA-seq on gastrocnemius muscle, liver, and white adipose tissue (38)), and *P. leucopus* dermal fibroblasts (qPCR, this study). *P. leucopus* consistently exhibits a higher male-to-female ratio than *M. musculus* across datasets. *M. musculus* qPCR samples included three biological replicates per sex: one embryonic fibroblast line and two adult ear-derived dermal fibroblast lines. All *P. leucopus* samples were ear-derived dermal fibroblasts from three biological replicates per sex. Bars represent the mean, and error bars indicates SEM. Statistical significance was assessed using one-way ANOVA (\*\*\*\**p* < 0.0001, ns = not significant). (D) RNA fluorescent *in situ* hybridization (FISH) using species-specific *Xist* probes (green) detects *Xist* RNA in dermal fibroblasts from *M. musculus* and *P. leucopus*. In female cells of both species, *Xist* RNA appears as a “cloud”. In contrast, *Xist* RNA is not detected in male *M. musculus* dermal fibroblasts; however, small *Xist* puncta (arrows) are observed in the nuclei of male *P. leucopus* dermal fibroblasts. Nuclei are counterstained with DAPI (blue). Representative images are shown from parallel hybridization experiments with three biological replicates. Scale bars = 10 μm. (E) RNA FISH of *P. leucopus* dermal fibroblasts stained with DAPI (blue) and *Xist* RNA probes (green) shows robust *Xist* signal in female nuclei and punctate *Xist* signal in a subset of male nuclei. Quantification indicates >90% *Xist*-positive nuclei in females and 60–70% in males (n = 3 biological replicates per sex). (F) RNA FISH of *Peromyscus leucopus* male and female dermal fibroblast metaphase chromosome spreads stained with DAPI (gray) and probed for *Xist* RNA (green) shows *Xist* signal localized to the X chromosome in both sexes. Representative female (XX) and male (XY) karyotypes are shown. (G) Representative X chromosomes from male and female cells showing sex-specific *Xist* RNA localization patterns by RNA FISH on metaphase chromosome spreads, alongside a schematic of the genomic location of *Xist* on the *P. leucopus* X chromosome (left).

Quantitative RT-qPCR confirmed that *Xist* expression in male *P. leucopus* is substantially elevated relative to the basal levels observed in male *M. musculus* (Fig. 2B). This difference was further reflected in male-to-female expression ratios, which are markedly higher in *P. leucopus* than in *M. musculus* (Fig. 2C). The *M. musculus* data include a male embryonic fibroblast sample, which, consistent with adult dermal fibroblasts, also exhibited minimal *Xist* expression. To contextualize our RT-qPCR results, we analyzed publicly available bulk RNA-seq data from *M. musculus* (38), which includes sex-matched transcriptomes from tissues such as gastrocnemius muscle, liver, and white adipose tissue. Consistent with RT-qPCR data, these datasets show robust *Xist* expression in females and negligible expression in males. The concordance between RNA-seq and qPCR datasets validates our expression measurements and underscores the distinct presence of *Xist* expression in male *P. leucopus*.

To determine whether *Xist* transcription is accompanied by canonical *Xist* RNA activity, we examined its nuclear localization by RNA fluorescence *in situ* hybridization (FISH) using species-specific probes. In female fibroblasts of both species, *Xist* formed the characteristic *Xist* RNA “cloud” associated with chromosome-wide coating of the inactive X chromosome (Fig. 2D), consistent with its well-established role in spreading-mediated silencing. Although present in both species, the female *P. leucopus* signal appeared to be more diffuse than the compact nuclear cloud observed in female *M. musculus*. As expected, no detectable *Xist* signal was observed in male *M. musculus* fibroblasts. In contrast, male *P. leucopus* fibroblasts displayed discrete, punctate *Xist* signals in 60-70 % of nuclei (Fig. 2D–E).

To determine whether male *Xist* localizes to the X chromosome, we performed RNA FISH on metaphase chromosome spreads. In females, *Xist* signal spreads across nearly the entire X chromosome, consistent with its established role in X chromosome inactivation. In males, however, the signal is confined to a discrete locus that precisely corresponds to the *Xist* genomic location identified through sequencing-based mapping (Fig. 1F,G). These data confirm that male *Xist* is transcribed from the X chromosome.

The punctate pattern observed in male *P. leucopus* is distinct from the canonical cloud-like *Xist* RNA distribution. Individual foci are typically singular, compact, and smaller than those in female nuclei, indicating that *Xist* RNA accumulation is spatially restricted and does not spread across the X chromosome. This constrained localization suggests that *Xist* in male cells may nucleate local chromatin interactions without achieving the distribution required for chromosome-wide silencing. The reproducibility of this pattern across dermal fibroblasts derived from three independent males further supports that it represents a stable regulatory state in a substantial subset of the analyzed cells.

### Expression of *Xist* in Male and Female *P. leucopus* Dermal Fibroblasts is Associated with Incomplete X-to-Autosome Dosage Compensation

To define the functional consequences of *Xist* expression in *P. leucopus*, we analyzed bulk RNA-seq data for ear-derived primary dermal fibroblasts isolated from adult males and females of *P. leucopus* and *M. musculus* (37), as well as liver tissue (39) from both species and sexes. In canonical models, *Xist*-mediated XCI reduces X-linked transcription in females to equalize the monoallelic X transcription in males, while compensatory upregulation of the active X ensures balanced overall transcriptional output between the X chromosome and autosomes in both sexes (40). Deviations from this framework therefore provide insight into how *Xist* activity influences genome-wide transcriptional balance. The comparative transcriptomic analysis enabled us to investigate expression levels of *Xist* across species and sexes, and to assess XCI with the extent of the X-to-autosome dosage compensation in *P. leucopus* and *M. musculus*.

Consistent with canonical XCI, female dermal fibroblasts and liver from both *M. musculus* and *P. leucopus* exhibited robust *Xist* expression (Fig. 3A). In dermal fibroblasts, *Xist* transcript levels were comparable between females of the two species, indicating conserved transcriptional regulation of *Xist*. In contrast, *M. musculus* males exhibited negligible *Xist* expression, whereas male *P. leucopus* displayed consistent and detectable *Xist* expression at levels significantly higher than those observed in *M. musculus* (Fig. 3A). This pattern extended to the liver tissue, where male *P. leucopus* also exhibited reproducible *Xist* expression across samples. The persistence of *Xist* expression across tissues, together with independent validation by RT-PCR and RNA FISH (Fig. 2), indicates that male *P. leucopus* somatic cells maintain sustained *Xist* activity, raising the possibility that downstream chromatin regulatory mechanisms are engaged outside the canonical female-specific context.

**Figure 3.**
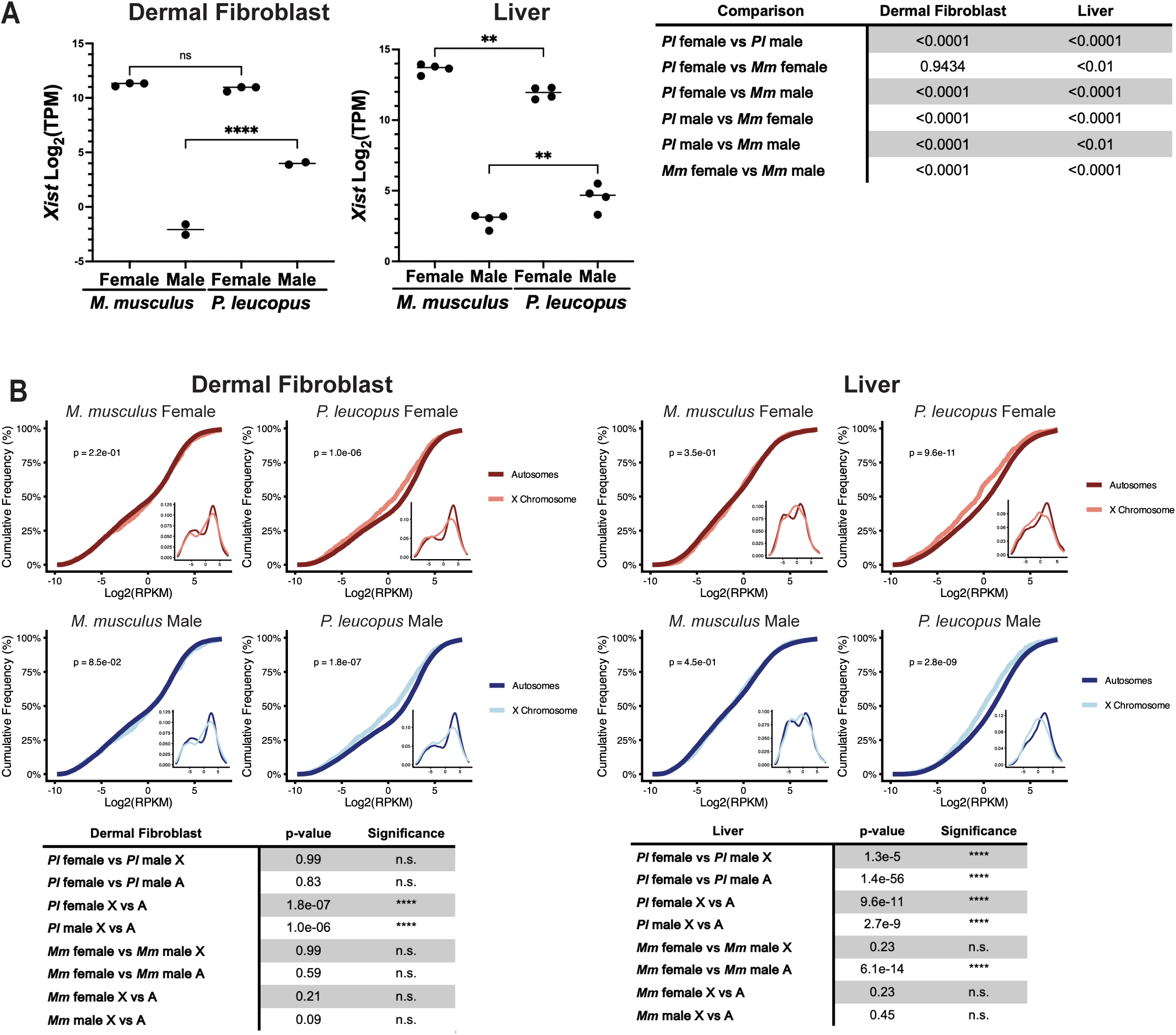
Consistent Male *Xist* expression and reduced X-linked dosage compensation in *P. leucopus*. (A) Bulk RNA-seq reveals detectable male *Xist* expression in *P. leucopus* in both primary cells (dermal fibroblasts) and tissue (liver). Statistical comparisons between species and sexes are provided in the accompanying tables (t-test). (B) Cumulative distribution plots (CDFs) of X-linked compared to autosomal gene expression (log RPKM) indicate complete X-to-autosome dosage compensation in *M. musculus*. In contrast, *P. leucopus* exhibits reduced X-linked gene expression relative to autosomes in both dermal fibroblasts and liver across sexes. CDFs are shown for each sample. The inset in each CDF panel shows the frequency of X-linked and autosomal gene expression levels for the corresponding sample. Kolmogorov-Smirnov test statistics are summarized in the table below.

To determine whether *Xist* expression is associated with dosage compensation, we compared X-linked and autosomal gene expression across sexes and tissues. Genes with zero expression (RPKM = 0) were excluded to focus on actively transcribed loci. Cumulative distribution analysis of logL-transformed RPKM values showed that in *M. musculus*, X-linked gene expression closely matched autosomal expression in both tissues and sexes, consistent with effective dosage compensation through XCI and X upregulation. In contrast, *P. leucopus* exhibited reduced X-linked expression relative to autosomes in both sexes and tissues, indicating a failure to achieve full X-to-autosome dosage compensation. This reduction in X-linked transcriptional output suggests that regulation of the *P. leucopus* X chromosome is not fully balanced with the presence of *Xist* activity.

To assess sex-specific effects, we further compared X-linked and autosomal gene expression between males and females. In *M. musculus*, X-linked gene expression was comparable between sexes in dermal fibroblasts, consistent with canonical XCI. In *P. leucopus*, however, liver tissue—but not dermal fibroblasts—showed sex-dependent differences in X-linked gene expression, indicating that *Xist* activity interacts with tissue-specific transcriptional programs. Autosomal genes also exhibited sex-biased expression in liver in both species, but not in dermal fibroblasts, suggesting that tissue-specific transcriptional variation contributes broadly to sex differences beyond the X chromosome.

Together, these results indicate that X-chromosome dosage compensation in *P. leucopus* differs from that in *M. musculus* and varies by tissue. Despite persistent *Xist* expression in both sexes, X-linked gene regulation in *P. leucopus* does not follow a uniform, chromosome-wide compensation program, but instead reflects a context-dependent regulatory state distinct from canonical XCI.

### *Xist* RNA in Female and Male *P. leucopus* Dermal Fibroblasts is Associated with Chromatin Silencing

XCI in eutherian mammals is mediated by the establishment of facultative heterochromatin, characterized by widespread deposition of H3K27me3 through Polycomb repressive complex 2 (PRC2) recruited by *Xist* RNA. To determine whether *Xist* expression in male *P. leucopus* is similarly associated with repressive chromatin, we performed combined RNA FISH for *Xist* and immunofluorescence (IF) for H3K27me3 in male and female *P. leucopus* dermal fibroblasts.

In female dermal fibroblasts, *Xist* RNA foci robustly colocalized with discrete H3K27me3-enriched domains, consistent with canonical XCI and *Xist*-mediated chromatin silencing (Fig. 4A, top row). In contrast, a subset of male dermal fibroblasts exhibited *Xist* RNA foci (Fig. 2D-E), a fraction of which colocalized with H3K27me3 (Fig. 4A, bottom row). Pixel intensity analysis of representative nuclei confirmed spatial overlap between *Xist* and H3K27me3 signals, with coincident peaks at the site of colocalization (Fig. 4B), indicating that *Xist* RNA can recruit repressive chromatin in male cells. This pattern was reproducibly observed in fibroblasts isolated from three independent male animals.

**Figure 4.**
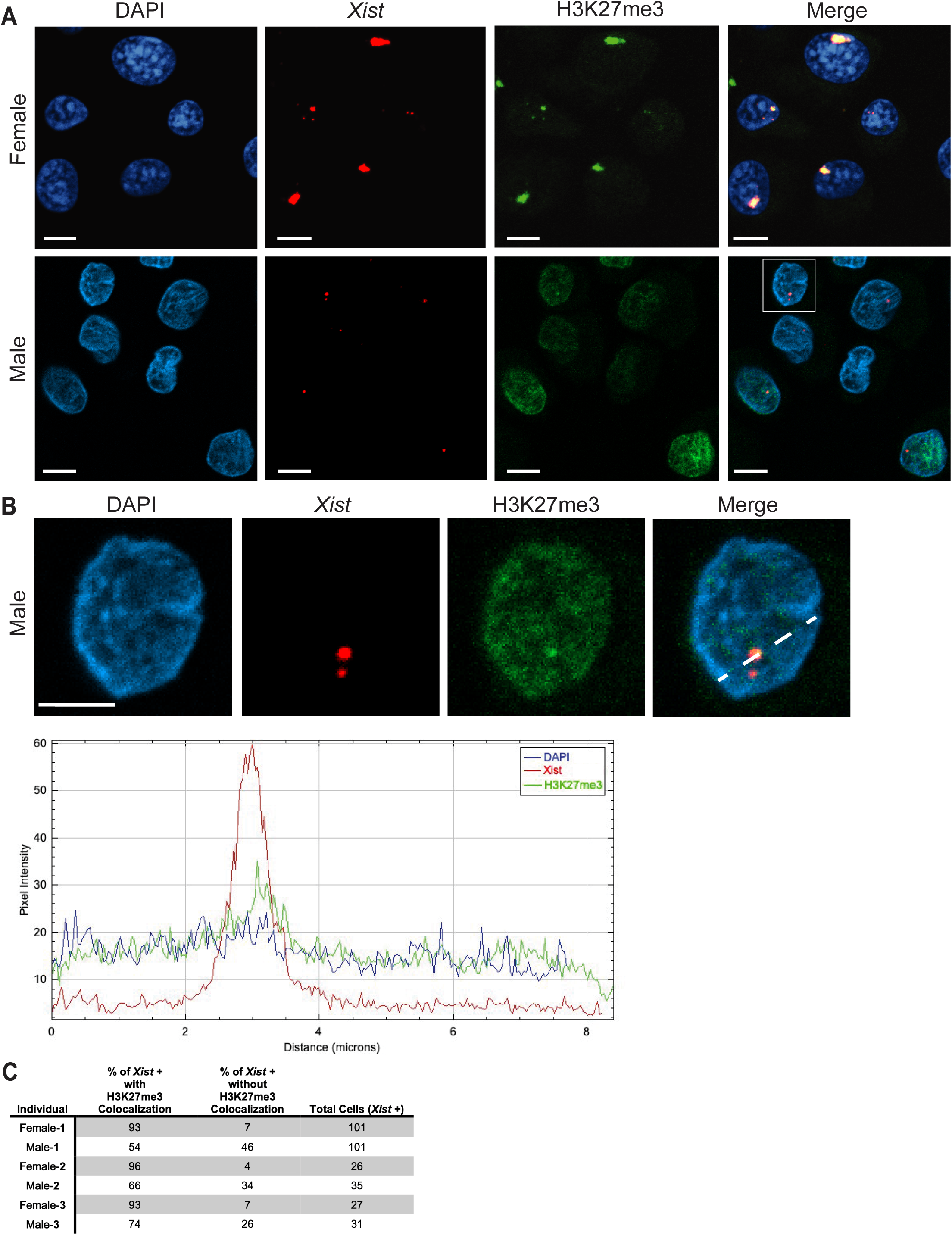
Male *Xist* is associated with deposition of H3K27me3 heterochromatin. (A) Immunofluorescence combined with RNA FISH analysis shows colocalization of H3K27me3 (green) and *Xist* RNA (red) in female and male *P. leucopus* dermal fibroblasts. DAPI (blue) marks cell nuclei. In female nuclei, *Xist* RNA colocalizes with H3K27me3, consistent with X-chromosome silencing. In a subset of male nuclei, *Xist* RNA also overlaps with H3K27me3. The white box indicates a representative male nucleus shown in Panel B. Images are shown as single Z-planes. Scale bars = 10 µm. (B) Representative image of a male *P. leucopus* cell showing colocalization of H3K27me3 (green) and *Xist* RNA (red) within the nucleus (DAPI, blue). A line profile plot along the indicated axis (white dashed line) quantifies signal intensities for DAPI, H3K27me3, and *Xist*, demonstrating spatial overlap of H3K27me3 and *Xist* signals. Scale bar = 10 µm. (C) Percentage of *Xist*-positive cells exhibiting colocalization of H3K27me3 and *Xist* RNA in *P. leucopus* dermal fibroblasts from three individual females or males.

However, the extent and frequency of *Xist*-H3K27me3 colocalization differed markedly between sexes. Only approximately 50% of *Xist*-positive male cells showed detectable colocalization, compared to more than 90% of female cells (Fig. 4C). These data indicate that, although *Xist* in male *P. leucopus* retains the capacity to recruit PRC2, this interaction is less frequent and likely less efficient than in the canonical female context. Reduced and spatially restricted PRC2 engagement would limit H3K27me3 deposition to discrete loci rather than establishing continuous, chromosome-wide silencing. As a result, repressive chromatin in male cells is likely weaker and less stable, insufficient to propagate across the X chromosome or enforce uniform silencing. Instead, *Xist*-mediated Polycomb recruitment is expected to remain localized, generating focal H3K27me3 enrichment that selectively represses nearby genes while leaving the majority of X-linked loci transcriptionally active.

Together, these findings indicate that *Xist* activity in male *P. leucopus* is distinct from its canonical spreading-based mechanism. Instead, sustained *Xist* transcription combined with spatially restricted RNA accumulation nucleates localized Polycomb-mediated repression but fails to achieve the spatial distribution required for chromosome-wide X chromosome silencing. This partial and heterogenous recruitment of PRC2 provides a mechanistic basis for the limited and gene-selective repression of X-linked genes in the male.

### H3K27me3 Reveals X-Linked Repression and Selective Target Genes in *P. leucopus*

The nuclear colocalization of *Xist* RNA with H3K27me3 (Fig. 4) prompted us to test whether *Xist* expression in *P. leucopus* males is associated with large-scale redistribution of repressive chromatin, and whether such redistribution is preferentially enriched on the X chromosome. To address this, we performed CUT&Tag profiling, a targeted chromatin profiling method that maps histone modifications genome-wide using antibody-directed tagmentation (41), for H3K27me3 enrichment in male and female *P. leucopus* and *M. musculus* dermal fibroblasts. To quantify chromatin distribution across chromosomes, we compared cumulative frequency distributions of H3K27me3 signal between the X chromosome and autosomes across species and sexes (Fig. 5A). This analysis revealed pronounced differences in X-linked H3K27me3 enrichment between *P. leucopus* and *M. musculus*, whereas autosomal distributions were relatively similar between species and comparable between sexes within each species. Notably, both female and male *P. leucopus* showed distinct X-chromosome enrichment relative to *M. musculus*, indicating a species-specific redistribution of repressive chromatin on the X.

**Figure 5.**
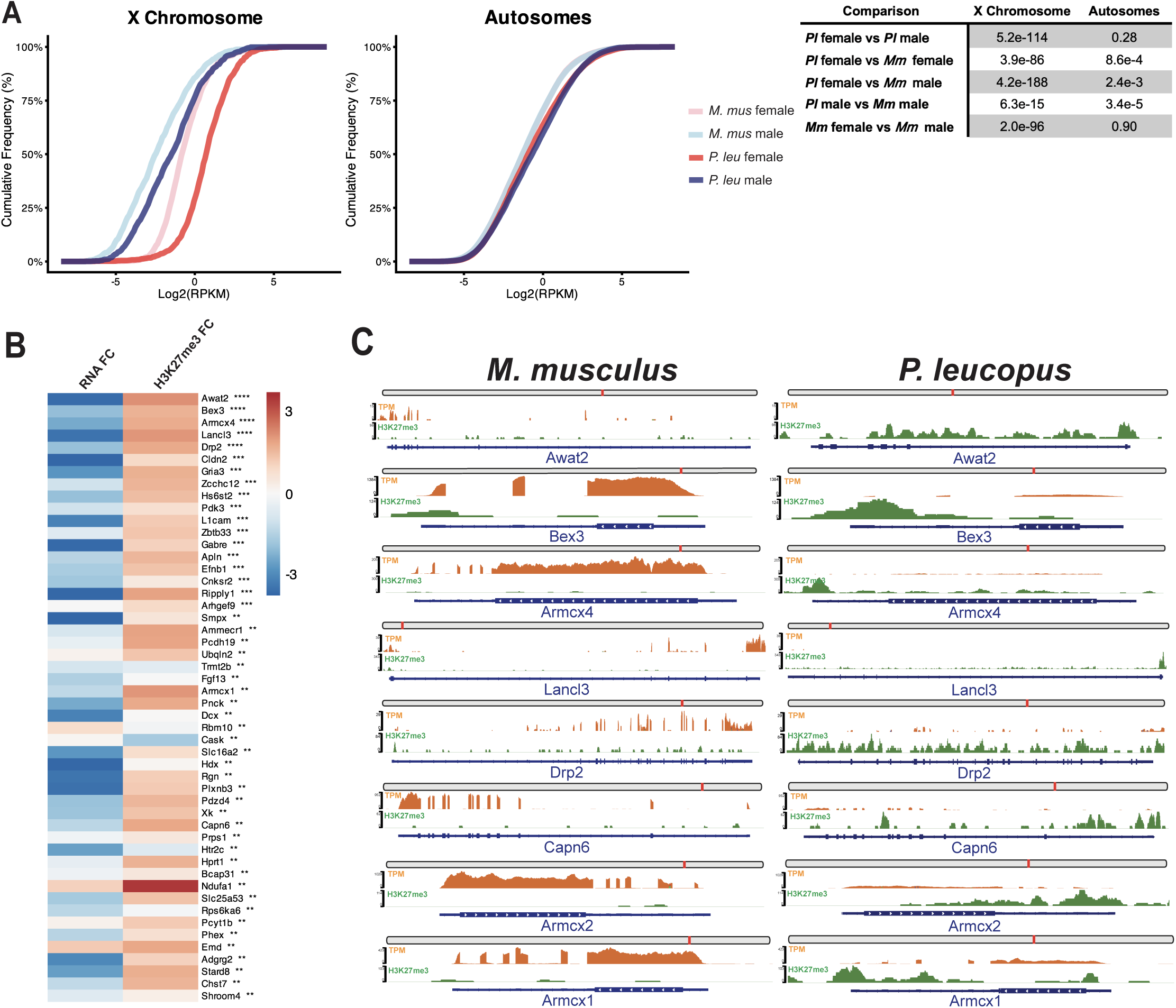
X-linked H3K27me3 enrichment and candidate gene silencing in *P. leucopus*. (A) CDF plot of H3K27me3 enrichment across the X chromosome and autosomes in male and females of both species. In *P. leucopus*, X chromosomes are right-shifted relative to autosomes in both sexes compared to *M. musculus*, indicating higher H3K27me3 enrichment. Kolmogorov-Smirnov test *p*-values and statistical significance for the indicated comparisons are provided in the table on the right. (B) Comparison of X-linked gene expression and H3K27me3 enrichment between *P. leucopus* and *M. musculus* males (dermal fibroblasts). Heatmap showing gene expression fold change (RNA FC) and H3K27me3 enrichment fold change (H3K27me3 FC) for the top 50 silenced X-linked genes, ranked by combined *p-value* derived from RNA FC and H3K27me3 FC between species. (C) Gene expression (TPM; orange) and H3K27me3 signal (CPM; green) across representative X-linked loci exhibiting strongest gene silencing in male *P. leucopus* (*Awat2, Bex3, Armcx4, Lancl3, Drp2, Capn6, Armcx2,* and *Armcx1*), shown along the gene body with corresponding genomic locations on the X chromosome.

Consistent with canonical XCI, both species exhibited clear female-male differences in X-linked H3K27me3 enrichment, with no corresponding differences on autosomes. However, in *P. leucopus*, the magnitude and distribution of X-linked H3K27me3 enrichment differed from *M. musculus*, suggesting a distinct mode of X-linked chromatin regulation. Importantly, the strong enrichment of H3K27me3 on the X chromosome, which far exceeds that on any the autosomes, provides chromatin-level evidence that the *P. leucopus* X chromosome is selectively repressed. This X-biased deposition of repressive marks is consistent with the reduced X-linked transcriptional output observed in *P. leucopus* (Fig. 3) and supports a model in which *Xist*-associated chromatin repression contributes to incomplete X-to-autosome dosage compensation.

Given the punctate nuclear distribution of *Xist* and H3K27me3 in *P. leucopus* males (Fig. 4C), we next tested whether X-linked H3K27me3 enrichment corresponds to gene-specific repression. Integrating CUT&Tag and RNA-seq datasets identified X-linked genes with elevated H3K27me3 and reduced expression in *P. leucopus* relative to *M. musculus*. Ranking genes by a combined p-value derived from both chromatin and expression fold-change differences yielded a set of candidate targets of selective repression (Fig. 5B), with *Awat2*, *Bex3*, *Armcx4*, *Lancl3*, and *Drp2* among the most silenced (Fig. 5C). Silenced genes included synaptic and axonal regulators *Drp2* and *Bex3* (42–45). Multiple members of the *Armcx* family (*Armcx1*, *Armcx2*, *Armcx4*) were co-repressed. This family is highly expressed in the nervous system and regulate neuronal function and mitochondrial trafficking, suggesting local chromatin architecture may facilitate silencing within this gene cluster (46).

To directly link gene repression to *Xist* activity, we performed Paired-Tag, a single-nucleus multi-omics approach that simultaneously profiles histone modifications and transcriptomes within the same cell, enabling direct coupling of chromatin state with gene expression at single-cell resolution (47). In *P. leucopus* males, *Xist*-positive nuclei exhibited increased H3K27me3 and reduced transcription at a subset of X-linked genes, including *Lancl3, Capn6, Armcx1,* and *Armcx2* (Supplementary Fig. 2). These genes were also silenced in *Xist*-positive female *P. leucopus* and *M. musculus* nuclei, but not in male *M. musculus*, further suggesting these genes are specific to male *P. leucopus Xist* silencing.

### *Xist* Expression is Coupled to X-linked Chromatin Repression

To determine whether the relationship between *Xist* expression, chromatin state, and transcriptional output represents a general property of *Xist*-positive nuclei, we compared H3K27me3 enrichment and X-linked gene expression between *Xist*-positive and *Xist*-negative nuclei across species and sexes (Fig. 6). Across all samples examined, *Xist*-positive nuclei exhibited increased H3K27me3 enrichment on the X chromosome accompanied by reduced X-linked transcription relative to *Xist*-negative nuclei. In contrast, autosomal gene expression and H3H27me3 levels were largely unchanged between *Xist*-positive and *Xist*-negative nuclei, indicating that *Xist*-associated chromatin and transcriptional effects are highly specific to the X chromosome.

**Figure 6.**
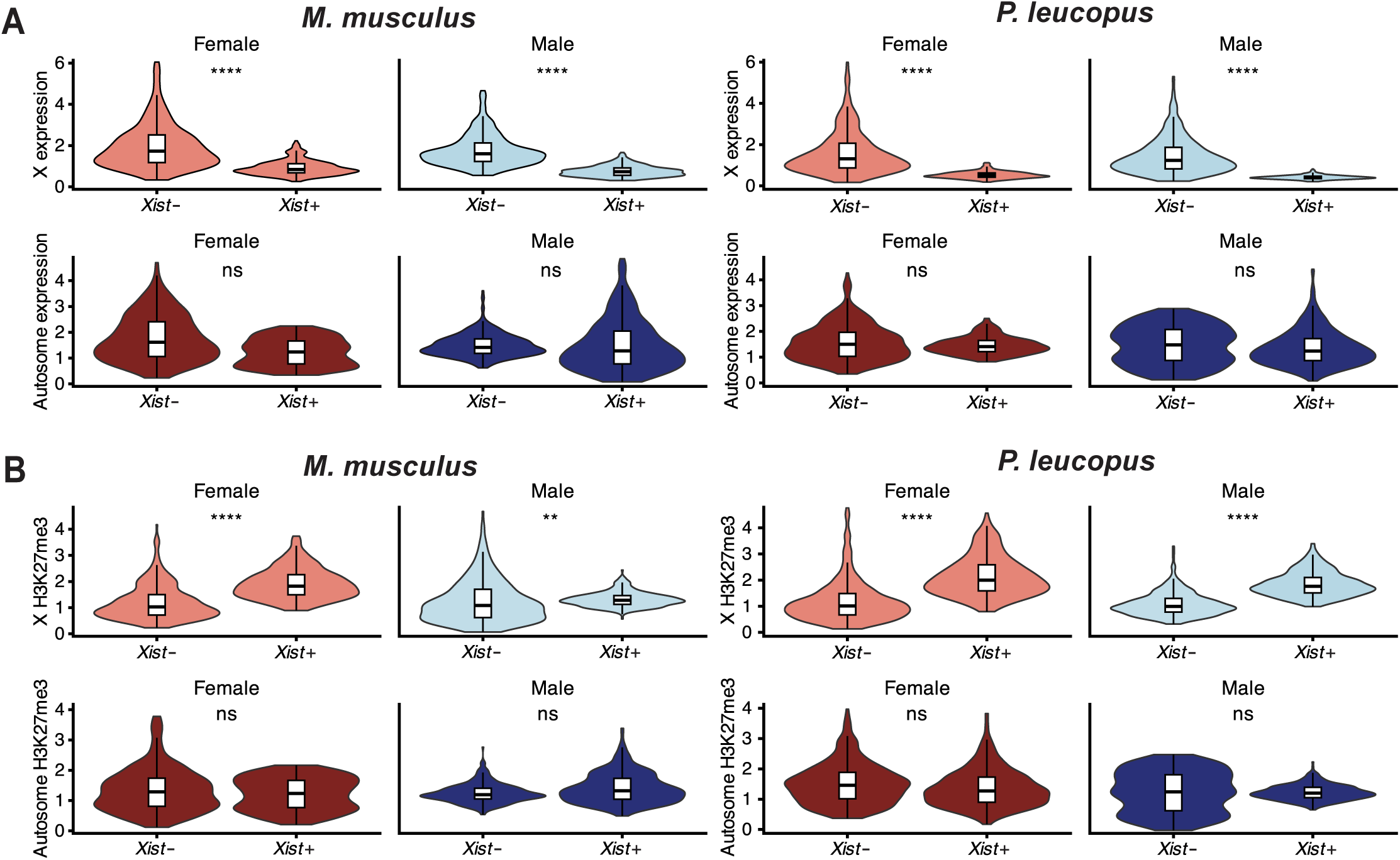
*Xist*-positive nuclei have reduced X-linked transcription and elevated H3K27me3. (A) Distribution of gene expression (log-normalized) for X-linked genes (top) and autosomes (bottom) in female (left) and male (right) *M. musculus* and *P. leucopus* liver cells, stratified by *Xist*-positive and *Xist*-negative nuclei. Statistical significance was assessed using a Wilcoxon rank-sum test. Significance levels are indicated as follows: ns (not significant), \**P* < 0.05, \*\**P* < 0.01, \*\*\**P* < 0.001, \*\*\*\**P* < 0.0001. (B) Distribution of H3K27me3 signal (log_2_CPM) for X-linked genes (top) and autosomes (bottom) in female (left) and male (right) *M. musculus* and *P. leucopus* liver cells, stratified by *Xist*-positive and *Xist*-negative nuclei. Statistical significance was assessed using a Wilcoxon rank-sum test with the same thresholds as in (A).

This pattern was observed consistently in both females and males of *P. leucopus*, as well as in female *M. musculus*, indicating that coupling between *Xist* expression and Polycomb-associated repression is a conserved feature of *Xist*-expressing cells. Importantly, a small population of *Xist*-positive nuclei was also detected in male *M. musculus*. These nuclei displayed the same chromatin and transcriptional signature—elevated H3K27me3 and reduced X-linked expression—as *Xist*-positive nuclei in female cells and in *P. leucopus* males. This finding indicates that the capacity for *Xist*-associated Polycomb recruitment and transcriptional repression is not restricted to female cells, but instead represents a cell-intrinsic regulatory potential that can be engaged across diverse biological contexts.

Together these results demonstrate that *Xist* expression is consistently associated with localized Polycomb-mediated repression at the single-cell level in male *P. leucopus*. However, in contrast to canonical chromosome-wide XCI, this coupling does not produce uniform silencing across the X chromosome. Instead, *Xist* functions within a context-dependent, gene-selective regulatory framework, generating spatially restricted repression that modulates subsets of X-linked genes.

## Discussion

X-chromosome inactivation (XCI) has long been conceptualized as a sequential and hierarchical process: *Xist* is expressed, spreads *in cis* across the X chromosome (48,49), recruits Polycomb repressive complexes (50,51), and enforces transcriptional silencing across most X-linked genes (52,53). This framework, derived primarily from studies in mice (3,6) and humans (5,54) has underpinned prevailing models of dosage compensation, in which *Xist* activity is coupled to chromosome-wide silencing. However, emerging evidence from marsupials (55) and monotremes (56) suggests that mammalian XCI strategies are more diverse than previously appreciated. Our findings in *P. leucopus* challenge this canonical paradigm. In a rodent with a conventional XY karyotype, we show that *Xist* is expressed in male somatic cells, associates with H3K27me3, and selectively represses a subset of X-linked genes, yet does not achieve chromosome-wide silencing or full X-to-autosome dosage compensation. These observations distinct *Xist* activity from its canonical outcome and suggests that its functional consequences are highly context-dependent.

A key question raised by these findings is how male *P. leucopus* tolerates *Xist* expression without deleterious effects. In *M. musculus,* ectopic *Xist* in XY cells induces chromosome-wide silencing of the single X chromosome and results in cell lethality (52,57), highlighting the incompatibility of canonical XCI with the male genomic context. Several features of the *P. leucopus* system likely mitigate this outcome. First, *Xist* accumulates at lower levels and localizes as discrete nuclear foci rather than spreading across the chromosome, indicating that its activity is spatially constrained from the outset. Consistent with this, *Xist*-H3K27me3 colocalization is detected in only a subset of the *Xist*-positive male nuclei, suggesting inefficient or heterogenous Polycomb engagement. Second, X-linked gene expression in *P. leucopus* is reduced relative to autosomes in both sexes, indicating incomplete transcriptional upregulation of the X chromosome. Under these conditions, *Xist* operates on a partially active chromatin substrate, such that additional local repression has limited global consequences. This regulatory architecture resembles X-linked dampening states observed prior to *Xist* upregulation in differentiating embryonic stem cells (58), raising the possibility that gene-selective repression represents a more ancestral or basal mode of X-chromosome regulation. In this framework, chromosome-wide silencing may be viewed as a specialized amplification of a more primitive locus-restricted regulatory capacity.

Persistent *Xist* expression in XY males is exceptionally rare among placental mammals, suggesting that the regulatory context in *P. leucopus* differs from canonical systems. In the creeping vole (*Microtus oregoni*), *Xist* expression occurs in the context of an atypical XX male karyotype, while monotremes lack *Xist* entirely and instead achieve partial dosage compensation through alternative mechanisms (56,59). Marsupials employes a distinct lncRNA, *Rsx* (*R*NA-on-the-*s*ilent *X*), which mediates incomplete and heterogeneous silencing (60,61). *P. leucopus* therefore represents a unique instance in which full-length *Xist* is expressed in XY somatic cells under non-pathological conditions. Notably, recent reports of *XIST* expression in male human peripheral glia (62) and cardiac pseudo-glial cells (63) further suggest that low-level, context-dependent *Xist* activity in male somatic cells may not be an isolated phenomenon. Together, these observations point to a broader, previously underappreciated capacity for context-dependent regulation by *Xist* across mammalian context.

At the gene level, our multi-omic analyses identify a consistent set of X-linked loci that are selectively repressed in *Xist*-positive male *P. leucopus* cells, including *Armcx1, Armcx2, Capn6,* and *Lancl3*. These genes are enriched in pathways governing mitochondrial dynamics, cytoskeletal organization, and metabolic regulation. *Armcx1* and *Armcx2* regulate mitochondrial transport and distribution, processes sensitive to gene dosage in neurons. Genes in the *Lancl* family are linked to metabolic stress responses, and *Capn6* plays a role in cytoskeletal remodeling as well as immune response (64–66). Importantly, these loci are not repressed in male *M. musculus*, indicating that *Xist* target selection is species-specific and context-dependent. Chromatin profiling further reveals that repression of these genes is associated with localized H3K27me3 enrichment, consistent with targeted Polycomb recruitment rather than chromosome-wide spreading. These findings support a model in which *Xist* functions as a selective modulator of gene expression, preferentially targeting dosage-sensitive loci rather than enforcing broad transcriptional silencing. In this view, *Xist* activity may be tuned to cellular context, fine-tuning gene expression in pathways where precise dosage control is critical.

An equally important question is how *P. leucopus* tolerates incomplete X-to-autosome dosage compensation. The Ohno hypothesis posits that XCI in females is coupled with upregulation of the active X chromosome in both sexes to equalize X-linked output relative to autosomes. While this model holds in mice (40,67) and humans (54), it is not observed in *P. leucopus*, where X:A expression ratios remain below 1.0 in both sexes across tissues. This persistent imbalance, together with elevated H3K27me3 enrichment on the X chromosome, indicates that *Xist-*associated repression contributes to a globally reduced transcriptional output from the X. The maintenance of organismal viability under these conditions suggests that strict dosage compensation is not a universal requirement in eutherian mammals. Instead, dosage compensation may exist along a continuum, with species-specific thresholds for tolerable imbalance. Although incomplete dosage compensation has been reported in platypus (56), birds (68,69), and certain insects (70), it has not previously been demonstrated in eutherian mammals. The ability of a placental mammal to remain viable without achieving full X-to-autosome balance challenges the assumption that tight dosage compensation is a universal feature of the eutherian lineage and instead suggests that the degree of X-chromosome compensation is more evolutionarily flexible than the canonical model implies.

Together, our results establish *Peromyscus leucopus* as a natural system in which *Xist*-mediated chromatin regulation is functionally decoupled from chromosome-wide XCI. Persistent *Xist* expression in males, its partial and spatially restricted association with Polycomb repression, and the selective repression of specific X-linked genes demonstrate that *Xist* can function as a context-dependent chromatin regulator rather than an obligate trigger of chromosome-wide silencing (71). More broadly, these findings suggest that *Xist* functions not only as a driver of chromosome-wide inactivation but also as a recruiter of chromatin-modifying machinery that can be deployed across varying spatial scales and regulatory contexts.

These findings have important implications for sex chromosome regulation and human disease. Variability in *XIST* expression, spreading, and escape from XCI is known to contribute to phenotypic heterogeneity in females (72,73), and our results suggest that intrinsic variability in *Xist* function may be a fundamental property rather than an exception. Therapeutic strategies that aim to harness *XIST* for chromosome-wide silencing (74,75) may therefore face intrinsic constraints, as *Xist*-mediated repression appears to depend on locus-specific features and chromatin context. More broadly, these findings highlight the evolutionary plasticity of lncRNA function and underscore the importance of comparative systems for uncovering general principles of chromosome regulation. Future studies across *Peromyscus* species, which exhibit diverse sex chromosome systems (76), may further elucidate the evolutionary pressures shaping dosage compensation strategies and identify the molecular determinants of *Xist* functional diversification.

## Methods

### Phylogenetic Trees

Whole-genome phylogenetic trees were constructed using PhyloT from NCBI taxonomy IDs: *P. leucopus* 10041; *P. maniculatus* 10042; *H. sapiens* 9606; *B. taurus* 9913; *M. oregoni* 111838; *M. musculus* 10090; *R. norvegicus* 10116; *O. cuniculus* 9986; *C. grisues* 10029. Trees were visualized with the interactive Tree of Life (iTOL)(77). *Xist* Repeats were analyzed using Tandem Repeat Finder (78) and adjusted manually. Repeat B was aligned using JalView and MAFFT default settings (79,80).

Phylogenetic analysis of *Xist* sequences was conducted using Clustal Omega (81,82) and the tree was generated using MEGA11 software (83). *Xist* sequences were aligned with available sequences from the NCBI accession number: *P. leucopus* XR_003736827.2; P. maniculatus XR_443108.3; H. sapiens NR_001564.3; B. taurus XR_001495594.3; M. oregoni XR_005973610.1; M. musculus NR_001463.3; R. norvegicus NR_132635.2; O. cuniculus XR_007912058.1; C. grisues XM_027432927.2.

### Cell Culture

*P. leucopus* and *M. musculus* dermal fibroblasts were derived from outbred LL stock maintained at the Peromyscus Genetic Stock Center (University of South Carolina), and *M. musculus* dermal fibroblasts were derived from outbred CD-1 stock obtained from Charles River Laboratories. Animals were bred at the UCI vivarium under protocols AUP-23-127 and AUP-24-042 approved by the UCI Institutional Animal Care and Use Committee (IACUC), and dermal fibroblasts were isolated from ear tissue using collagenase digestion and standard culture procedures as previously described (37).

Cells were cultured in Roswell Park Memorial Institute (RPMI) 1640 medium supplemented with 10% fetal bovine serum (FBS), 2 mM glutamine, 100 μM asparagine, 1% penicillin-streptomycin solution. Primary *M. musculus* dermal fibroblasts were isolated from E13.5 DR4 embryos after removal of brain and visceral tissue. The cells were cultured in Dulbecco’s Modified Eagle Medium (DMEM) supplemented with 10% FBS and 1% penicillin-streptomycin solution. Cells were grown at 37°C in 5% CO2. Primary dermal fibroblasts of low-passage cultures were used throughout this study.

### Metaphase Spread and Chromosome Analysis

Metaphase spreads were prepared from actively dividing fibroblasts treated with KaryoMAX colcemid solution (0.1 μg/mL, 2-4 h) (Thermo Fisher). Cells were harvested, incubated in hypotonic KCl (0.075 M, 15 min, 37°C), and fixed in Carnoy’s fixative (3:1 methanol:acetic acid). Fixed cells were dropped onto slides and air-dried overnight. Chromosomes were stained with 2% Giemsa stain (for chromosome counting) or RNA FISH was performed on the slides as described below (for visualizing *Xist* localization on the chromosomes).

### Reverse Transcription Quantitative PCR (RT-qPCR) and DNA Gel Electrophoresis

RT-PCR was performed to assess *Xist* transcript presence and to verify cDNA quality in all samples. Primers targeting *P. leucopus Xist* exon 1 were: PL-XIST-F1-qPCR (5′-TGAGACACTTTGCTGAACCC-3′) and PL-XIST-R1-qPCR (5′- GGGAGGCTTTGAGAACTAGATG-3′). Primers targeting *P. leucopus Gapdh* (PL-GAPDH-F1: 5′-TCACCACCATGGAGAAGGC-3′; PL-GAPDH-R1: 5′- GCTAAGCAGTTGGTGGTGCA-3′) were designed based on the *P. leucopus* genome annotation. Total RNA was extracted from male and female *M. musculus* and *P. leucopus* primary dermal fibroblasts using TRIzol and chloroform. cDNA was generated from total RNA using Superscript III RT (LifeTechnologies) for reverse transcription. *Xist* qPCR was performed using iSYBR Green (BioRad) on a BioRad Real-Time PCR System. The PCR products were run on a 2% agarose gel and visualized with Ethidium Bromide to verify that primers were amplifying the reverse transcript cDNA product.

### RNA Fluorescence In-Situ Hybridization (FISH)

RNA FISH was performed to observe the localization of *Xist* RNA in *M. musculus* and *P. leucopus* male and female primary dermal fibroblasts. The cells were fixed with 4% paraformaldehyde onto microscope slides. Cells were hybridized with *Xist* RNA probes specific to each species following the RNAscope Multiplex Fluorescent Assay v2 (Advanced Cell Diagnostics) protocol. Images were captured using a Carl Zeiss LSM 900 with Airyscan 2 confocal microscope and analyzed on Imaris software.

### Immunofluorescence

Following RNA FISH, fixed cells were blocked with PBS, 0.2% Tween20, 1% BSA. Anti-H3K27me3 (no. 39535; Active Motif) (1:100), and incubated at room temperature for 1 hour, then washed in PBS and 0.2% Tween20 3x for 5 min, were incubated with secondary antibodies at 1:500 for 1 hour at room temperature followed by three more washings and were mounted in VECTASHIELD with DAPI (Vector Laboratories). Images were captured using a Carl Zeiss LSM 900 with Airyscan 2 confocal microscope and analyzed on Imaris software.

Colocalization between *Xist* RNA FISH signals and H3K27me3 immunofluorescence was quantified using ImageJ (NIH). Pixel intensity values were measured across individual nuclei, and colocalization was assessed by comparing the overlap of fluorescence signals within defined regions of interest.

### RNA Sequencing and Data Analysis

As described in Duong et al. (37), total RNA was extracted with the Quick-RNA Miniprep Plus Kit (Zymo Research) including on-column DNase treatment, and cDNA libraries were prepared using the Illumina TruSeq Stranded mRNA kit. Multiplexed libraries were sequenced on an Illumina NovaSeq 6000 platform (150 bp paired-end; ∼120–250 million reads/sample). Raw RNA sequencing data are available under the GEO accession (GSM9100090, GSM9100093, GSM9100096, GSM9100099, GSM9100102, GSM9100069, GSM9100072, GSM9100075, GSM9100078, and GSM9100081).

Raw sequencing reads were trimmed to remove adapter sequences and low-quality bases, then analyzed using CLC Genomics Workbench (Qiagen) following the pipeline described in Duong et al. (37). Reads were aligned to the *Mus musculus* (GRCm39) and *Peromyscus leucopus* LL stock (GCF_004664715.2) coding sequences with a similarity fraction of 0.9, length fraction of 0.35, and a cost of 3 for mismatches, insertions, and deletions. Expression levels were reported in TPM and RPKM.

Cumulative distribution functions (CDFs) were generated separately for X-linked and autosomal genes in males and females. Comparisons between expression distributions were performed using two-sample Kolmogorov–Smirnov (KS) tests. All analyses and plotting were conducted in R using the dplyr, ggplot2, patchwork, and scales packages.

For male to female *Xist* expression ratio in *M. musculus*, we analyzed bulk RNA-seq data from a publicly available sex-matched dataset (38): Count and metadata files were obtained from the authors’ GitHub repository, and expression values were calculated and analyzed following the published data processing pipeline. To compare X chromosome expression relative to autosomes, we calculated X:A ratios using logL-transformed RPKM values. Raw RPKM values were averaged across male and female samples within each species, and a pseudocount of 0.0001 was added before log2 transformation to avoid undefined values. Genes with zero expression across all samples were excluded to remove sex-biased genes. Chromosome assignments were based on the *P. leucopus* reference genome (GCF_004664715.2), and genes were classified as either X-linked or autosomal.

### CUT&Tag Analysis

Bulk CUT&Tag (Cleavage Under Targets and Tagmentation) was performed to map H3K27me3 histone modifications across the genome in *M. musculus* and *P. leucopus* in primary dermal fibroblasts. Libraries were prepared by Epigenome Technologies, San Diego, CA. Raw sequencing reads were processed using their standard bioinformatics pipeline. Briefly, reads were trimmed for adapter sequences and low-quality bases using Trim Galore, then aligned to the mouse reference genome (GRCm39) for *M. musculus* samples and the deer mouse reference genome (GCF_004664715.2) for *P. leucopus* samples using Bowtie2 with default parameters. Duplicate reads were removed using Picard MarkDuplicates. Genome-wide coverage tracks were generated in BigWig format using deepTools bamCoverage with RPKM (Reads Per Kilobase per Million mapped reads) normalization and a bin size of 5 Kb. Data quality was evaluated based on H3K27me3 enrichment at transcriptionally silent genes across samples (Supplementary Fig. 3).

### Single-nucleus Paired-Tag Analysis

Droplet-based single-nucleus Paired-Tag profiling of H3K27me3 and transcriptomes was performed by Epigenome Technologies. Liver samples from sexually mature male and female *P. leucopus* and *M. musculus* were collected. The animals were comparable in size and age; *P. leucopus* were approximately one year old, while *M. musculus* were 10 weeks old. All tissue collection procedures were conducted under IACUC approval (AUP-23-127). Sequencing data from RNA and DNA libraries were processed using 10x Genomics pipelines. RNA libraries were run through CellRanger v7.0 with the chemistry flag set to --chemistry ARC-v1. DNA libraries were processed with CellRanger-ATAC using --chemistry ARC-v1. Raw FASTQ files were input to each pipeline using GCF_004664715.2_UCI_PerLeu_2.1 (deermouse, built using CellRanger-ARC mkref) or the 10X-provided refdata-cellranger-arc-mm10-2020-A-2.0.0 reference. Standard CellRanger outputs (filtered feature-barcode matrices for RNA; fragment files and peak/barcode matrices for ATAC) were retained for downstream analysis.

Extended oligo counting was used to identify sample-level barcodes within each assay. Briefly, For each cellular barcode, counts of the sample-specific extended oligo sequences were tabulated. A sample assignment was made when a single extended oligo sequence represented >90% identity of the extended-oligo reads associated with that cellular barcode. Barcodes failing to meet this threshold were marked as ambiguous and excluded from sample-specific downstream analyses. Cross-assay barcode mapping between RNA and DNA modalities was performed using the 10x Chromium ARC-bc-mapping file. RNA and DNA barcodes were matched by the mapping file to link transcriptome and chromatin accessibility data for the same physical cell.

DNA fragments from the CellRanger-ATAC-produced H3K27me3 fragment file were quantified in two complementary ways. First, fragments were aggregated into fixed-size genomic bins to produce a genome-wide accessibility matrix. Second, fragments overlapping candidate regulatory regions were counted to produce a peak-by-cell fragment matrix. Binning used non-overlapping windows of size 5 kb unless otherwise noted. Fragment counting used exact overlap with bin or peak coordinates; fragments overlapping multiple features were counted for each overlapping feature.

Peaks were called on aggregated ATAC fragments using MACS2 with parameters optimized for broad accessibility regions (--broad --broad-cutoff 0.1 --q 0.1). Cells were retained for downstream analyses only if they satisfied both RNA and DNA quality thresholds and were not flagged as multiplets. Retention criteria were: RNA UMI count >= 750 and DNA fragment count >= 750, and not flagged as a multiplet/ambiguous by barcode identification (see above). Cells failing any of these criteria were excluded from subsequent analyses. Summary QC metrics including UMI counts, fragment counts, fraction of reads in peaks, and mitochondrial RNA fraction were computed for all retained cells.

Paired-Tag data were used to jointly profile RNA expression and H3K27me3 enrichment at single-cell resolution. Cells with *Xist* expression >0 were classified as *Xist*-positive, while cells lacking detectable *Xist* expression were classified as *Xist*-negative. Data were normalized either by CPM or by log-normalization using Seurat, depending on the analysis. Only orthologous genes shared between species were retained for downstream analyses, and genes with zero expression or zero H3K27me3 signal across all cells were excluded. X-linked and autosomal genes were analyzed separately to compare RNA expression and H3K27me3 enrichment between *Xist*-positive and *Xist*-negative cells. Silenced genes were defined as X-linked genes showing reduced RNA expression in *Xist*-positive compared to *Xist*-negative cells, together with increases in H3K27me3 enrichment. X-linked and autosomal genes were then analyzed separately to compare RNA expression and H3K27me3 patterns between conditions. Chromosome-level analyses were performed by aggregating normalized signals across genes on each chromosome.

## Data Availability

Raw Paired-Tag sequencing data generated in this study have been deposited in the NCBI Sequence Read Archive under BioProject accession PRJNA1466601. Bulk CUT&Tag sequencing data have been deposited under BioProject accession PRJNA1466562.

## Acknowledgements

We thank members of the Sun lab for helpful discussion and Dr. Adeela Syed for assistance with microscopy. This work was funded by the National Institutes of Health through the National Institute of General Medical Sciences award R01GM141424 (S.S.) and the National Institute of Allergy and Infectious Diseases award R01AI157513 (A.D.L and A.G.B.). M.D.L.A.L. received support from Dr. Star Lee and the U.S. Department of Education’s Graduate Assistance in Areas of National Need (GAANN) fellowship program. This study was made possible in part through access to the Optical Biology Core Facility of the Developmental Biology Center, a shared resource supported by the Cancer Center Support Grant (CA-62203).

---

**<u>Supplementary Figure 1. Distance Phylogeny of *Xist* within cricetids.</u>**

Distance phylogram of *Xist* nucleotide sequences within cricetids including *P. maniculatus*, *P. leucopus*, *P. californicus*, and *O. torridus*, with *M. musculus* serving as the outgroup; node labels indicate percent bootstrap support (left). Overall clustering among the same cricetid species based on whole-genome taxonomy (right).

**<u>Supplementary Figure 2. Expression and H3K27me3 enrichment of candidate genes associated with male *P. leucopus Xist* across species, sexes, and *Xist* status.</u>**

(A) Gene expression levels of candidate silenced genes, shown as log_2_ CPM in *M. musculus* and *P. leucopus* male and female liver cells, stratified by *Xist*-positive and *Xist*-negative nuclei.

(B) H3K27me3 enrichment for the same genes, shown as log_2_ CPM in *M. musculus* and *P. leucopus* male and female liver cells, stratified by *Xist*-positive and *Xist*-negative nuclei.

**<u>Supplementary Figure 3. H3K27me3 CUT&Tag quality control and enrichment patterns.</u>**

Heatmap of H3K27me3 CUT&Tag signal across the genome, with genes ranked by average RNA-seq expression (TPM; high to low), in four biological samples (female: F1, F2; male: M1, M2) for *M. musculus* (top) and *P. leucopus* (bottom). Profile plots (top panels) show mean H3K27me3 signal centered around transcriptional start sites (TSS) and transcriptional end sites (TES) (±2 kb). Signal intensity is normalized to reads per kilobase per million mapped reads (RPKM). H3K27me3 signal is enriched at transcriptionally silent or lowly expressed genes.

## Notes

### Competing Interest Statement

The authors have declared no competing interest.

### Summary of Updates

The author list was re-ordered to be accurate.

