## Supplementary Figures for "*Xist* expression in male *Peromyscus leucopus* is associated with restricted chromatin repression and incomplete X-to-autosome dosage compensation"

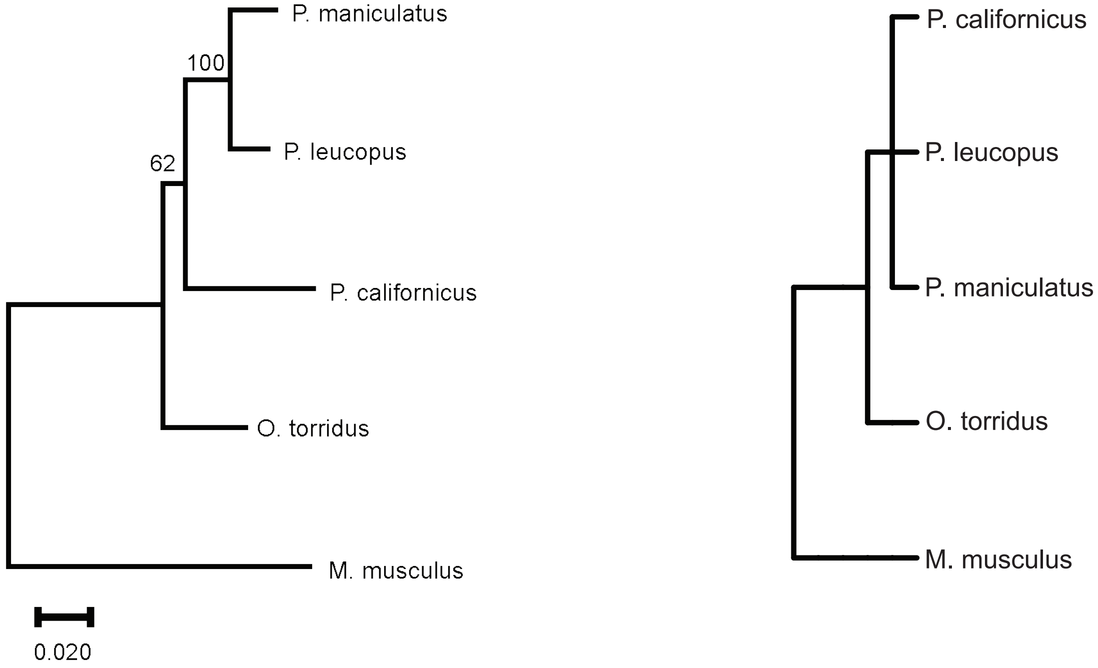


Supplementary Figure 1. Distance Phylogeny of *Xist* within cricetids.

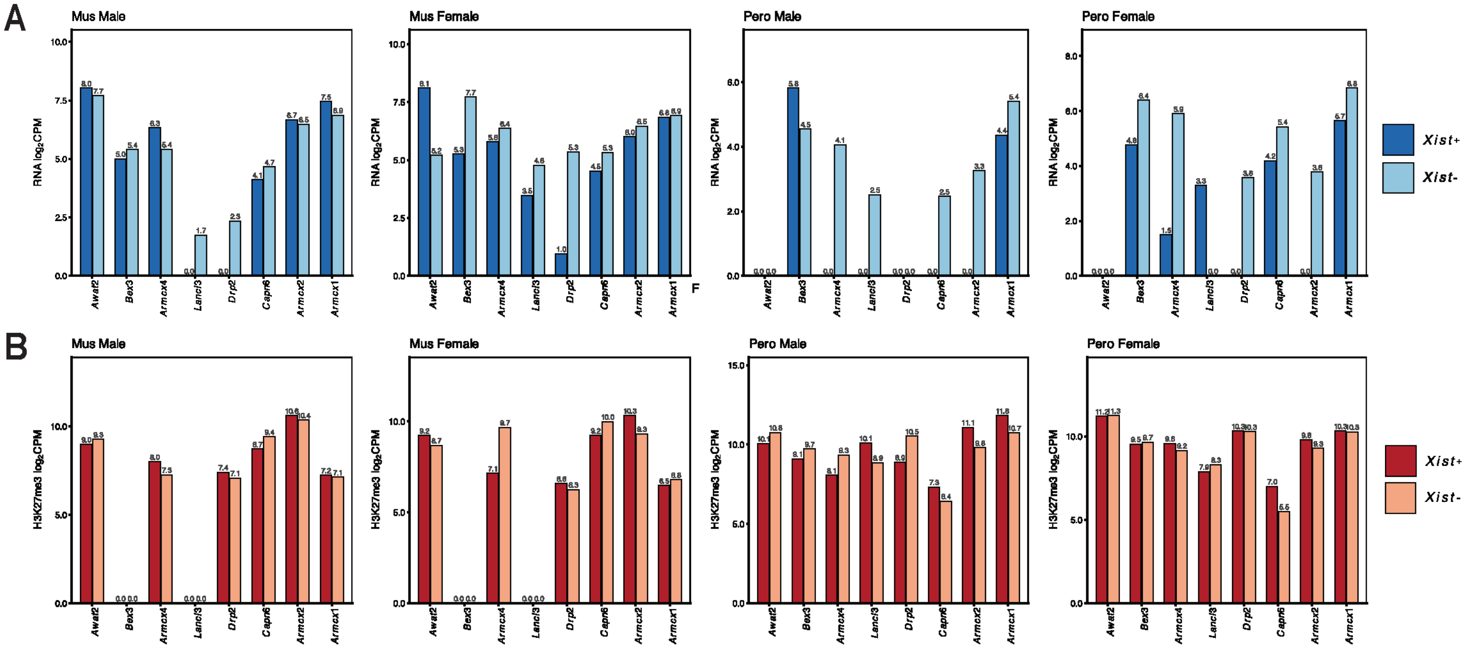


Supplementary Figure 2. Expression and H3K27me3 enrichment of candidate genes associated with male *P. leucopus Xist* across species, sexes, and *Xist* status.

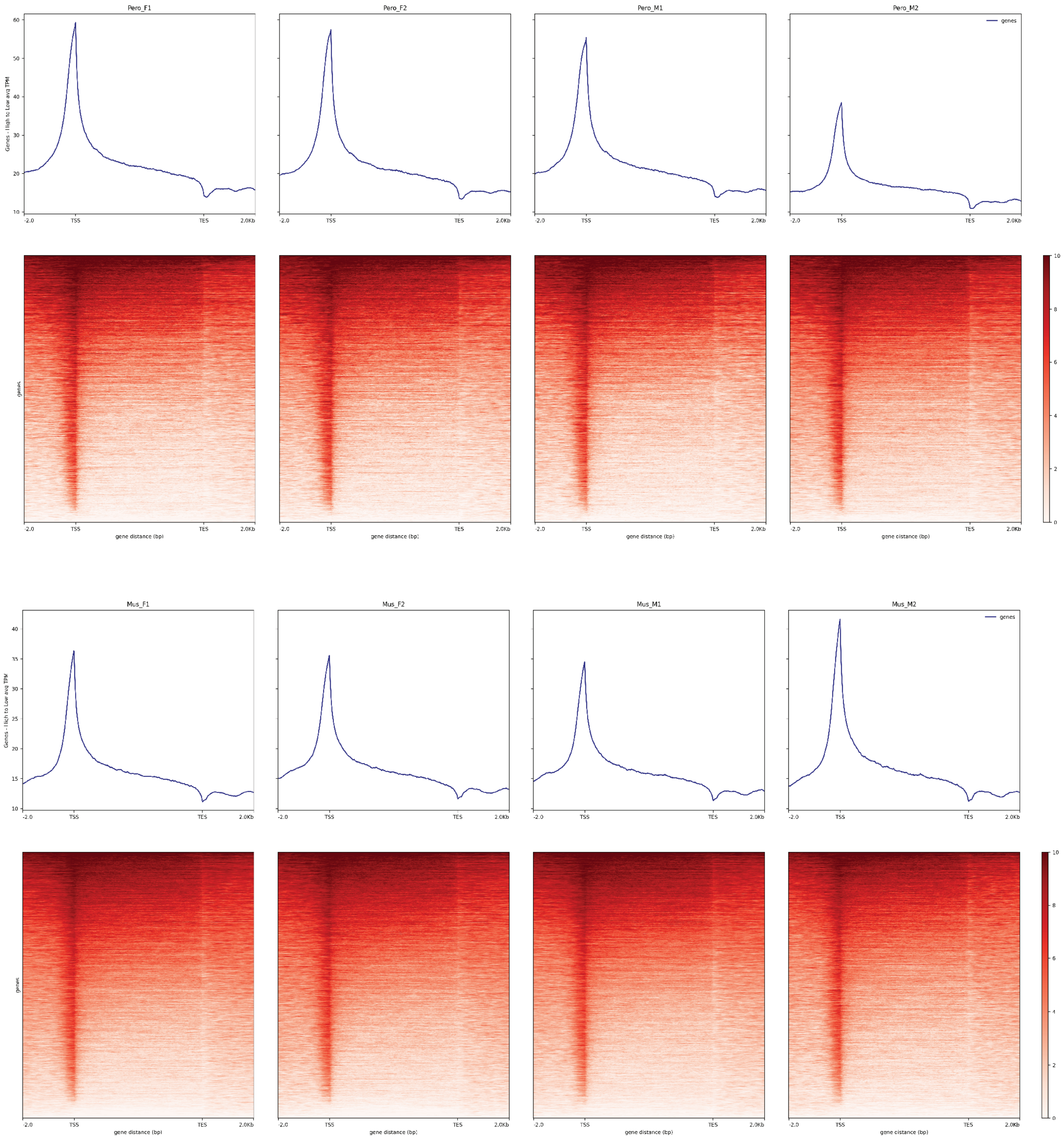


Supplementary Figure 3. H3K27me3 CUT&Tag quality control and enrichment patterns.
